# Multisensory interactions on auditory and somatosensory information in expert pianists

**DOI:** 10.1101/2022.02.23.481694

**Authors:** Masato Hirano, Shinichi Furuya

## Abstract

Fine-tuned sensory functions are bases of efficient motor control and learning and typically characterize skilled individuals. Although numerous studies demonstrated enhanced unimodal sensory functions at both neural and behavioral levels in skilled individuals, little is known about their multisensory interaction, especially multisensory integration and selective attention that involve volitional control of information derived from multiple sensory organs. Here, we show unique multisensory interaction functions of expert pianists. Expert pianists and musically untrained individuals performed five sets of intensity discrimination tasks at the auditory and somatosensory modalities with different conditions: (1) auditory stimulus, (2) somatosensory stimulus, (3) congruent auditory and somatosensory stimuli (i.e., multisensory integration), (4) auditory and task-irrelevant somatosensory stimuli, and (5) somatosensory and task-irrelevant auditory stimuli. In the fourth and fifth conditions, participants were instructed to ignore a task-irrelevant stimulus and to pay attention to a task-relevant stimulus (i.e., selective attention). The unimodal intensity discrimination of the pianists was superior to that of the nonmusicians at the auditory modality but not at the somatosensory modality. While the discrimination perception was superior in the condition (3) compared to the better one of the individual unimodal conditions (i.e., conditions 1 and 2) only in the pianists, the task-irrelevant somatosensory stimulus worsened the auditory discrimination more in the pianists than the nonmusicians. These findings indicate efficient processing of multisensory information in expert pianists, which enables to benefit from multisensory integration of the auditory and somatosensory information, but exacerbates top-down selective inhibition of somatosensory information during auditory processing.

## Introduction

We perceive the external world via interactions of multisensory information derived from different sensory organs and utilize the perceived information to update motor outputs that produce the desired movements. In the real world, our experiences have multisensory features. Integration of multisensory information (i.e. multisensory integration) shapes accurate perception when each modality of sensory information represents congruent features ^1^. For example, simultaneous presentation of visual and auditory stimuli enhances both signal detectability and motor reaction time on the detection of a spatial location from which the two sensory stimuli originate^2^. By contrast, in some cases such as noisy environments, attending to sensory signals of a particular modality increases signal detectability^3^, which accompanies ignoring task-irrelevant sensory information (i.e., selective attention). Previous studies demonstrated that these multisensory interaction functions are acquired at a late stage of development and are adapted with age^4,5^ and experiences^6^. This suggests that multisensory interaction functions have the potential of plastic adaptation through a course of abundant multisensory experiences.

Plastic changes in unimodal sensory functions have been investigated in previous studies of sensory and motor learning ^7,8^. Unimodal sensory functions can be sharpened with repeated exposure to sensory stimuli of a single modality or with motor training ^9–13^. Furthermore, previous studies demonstrated the plastic adaptation of multisensory interaction functions in trained individuals ^13–16^. A common approach among these studies is the manipulation of congruency of sensory information between multiple modalities. For example, expert musicians can more accurately perceive incongruent stimuli between auditory and visual sensory information, like musical score and sounds, compared with musically untrained individuals ^15,17,18^. However, it remains unclear whether the above-mentioned multisensory interaction functions (i.e., multisensory integration and selective attention) also benefit from extensive multisensory training. These multisensory interaction functions involve volitional control of information derived from the multisensory sensory organs (i.e., whether integrate or suppress), which is different from the incongruent stimuli detection that identifies differences in information encoded in sensory signals of different modalities. Recent studies demonstrated that multisensory integration involves combining multisensory sensory information by summing prior knowledge and the independent stimulus estimates from each modality ^19–21^. On the other hand, the selective attention in multisensory situations is defined as guiding attention toward modalities that represent task-relevant information and away from modalities that provide task-irrelevant information and/or noise. Previous studies demonstrated that short-term playing of an action video game improves visual selective attention that is characterized as guiding attention toward task-relevant visual stimuli and away from irrelevant visual disturbances ^22,23^. However, to the best of our knowledge, none of the studies examined plasticity of selective attention in multisensory situations, especially in the auditory and somatosensory modalities.

Expert musicians are a unique population to address these issues because playing musical instruments provides extensive auditory and somatosensory experiences. In piano performance, sensory inputs from the auditory and somatosensory modalities provide abundant information such as loudness and timbre of notes. Indeed, previous studies have demonstrated that such multisensory experiences provide pianists with superior perceptual abilities in both modalities over untrained healthy individuals^11,24,25^. On the other hand, previous studies demonstrated interactions of sensory perception between these two modalities^26–28^. Such auditory-somatosensory integration plays important roles in successful piano performance and thus would be plastically adapted through long-term piano training. One of the important requirements for expressive music performance is fine control of the intensity of sounds. The sound intensity is perceived mainly from the auditory modality, but also from the somatosensory modality that encodes information on the sound intensity via the keystroke (e.g., pressure sense at fingertips). In addition, accurate perception of the sound intensity necessitates both the integration and selective attention of information derived from these two modalities. Expert pianists thus have undergone unique multisensory experiences from the auditory and somatosensory modalities since childhood. This raises the hypothesis that expert pianists have superior multisensory interaction functions in the intensity domain between the auditory and somatosensory over the musically untrained individuals. Here, we designed a series of psychophysical experiments to probe plasticity of the multisensory interaction functions in the auditory and somatosensory modalities, especially with respect to the multisensory integration and selective attention, through a comparison between expert pianists and nonmusicians.

## Materials and methods

### Participants

Fifteen pianists (22.9±4.5 years old [mean±SD], 11 females) and 15 musically untrained individuals (nonmusicians: 25.9±3.9 years old, 6 females) participated in the present study. All pianists majored in piano performance in a musical conservatory and/or had extensive and continuous private piano training under the supervision of a professional pianist/piano professor. By contrast, all nonmusicians have absolutely no experience of training of musical instruments except for mandatory musical education program during the elementary school period. All participants gave their written informed consent before participating in the experiments. All experimental procedures were approved by the ethics committee of Sony Corporation.

### Procedure

We firstly assessed the detection threshold for each of the somatosensory and auditory stimuli by delivering the sensory stimulus at the pace of 0.25 to 0.5 Hz with decreasing/increasing the stimulus intensity. Participants were asked to press a key if they detect a sensory stimulus. The stimulus intensity was decreased/increased by 2 dB if the reaction time between the sensory stimulus and the keypress was below/over 500 ms. The experimenter stopped the task if the time course of the stimulus intensity reached a plateau. The detection threshold was defined as the averaged intensity across the last 10 trials. The detection threshold did not differ between the two groups (auditory: pianists: −40.62±4.88 dB (0 dB corresponds to the sound volume of 70 dB speaker level (dBSPL) just in front of the speaker); nonmusicians: −40.41±7.10 dB; unpaired t-test: t=-0.09, p=0.93; somatosensory: pianists: −17.19±3.38 dB (The vibration intensity when an AC voltage of 10 V is applied to the piezoelectric actuator is 0 dB); nonmusicians: −16.04±2.78 dB; unpaired t-test: t=-0.87, p=0.40).

After detecting the detection thresholds of the two modalities, participants were performed the intensity discrimination test 5 times with different conditions; (1) auditory condition (A), (2) vibrotactile condition (V), (3) multisensory integration condition (A+V: auditory + vibrotactile), (4) auditory selective attention condition (Aatt), and (5) vibrotactile selective attention condition (Vatt). The order of the conditions was randomized across participants. In the A+V condition, we delivered the auditory and vibrotactile stimuli simultaneously. The intensity of the comparing stimulus differed from that of the standard stimulus by the same level between the two modalities. In the Aatt condition, we delivered the vibrotactile and auditory stimuli simultaneously, and manipulated the intensity of the comparing stimulus in the auditory modality without changing the intensity of the comparing stimulus in the vibrotactile modality as that of the standard stimulus. Thus, the vibrotactile stimuli represented no information on the discrimination task (i.e. task-irrelevant information). The procedure of the Vatt condition was the reverse condition of the Aatt, indicating that the auditory stimuli had no information on the task. Both the auditory and vibrotactile stimuli were delivered via different channels of the same D/A converter, which allowed us to precisely synchronize the two stimuli. This synchronization was validated in our pilot experiment.

### Auditory stimulus

Each auditory stimulus consisted of 2000 Hz pure tone and lasted for 0.2 s. We used this sound frequency because human auditory perception is highly sensitive at this frequency^29^. In order to eliminate pop noise, the amplitude of the auditory stimulus linearly decreased to silent during the last 100 ms. The sound signal was produced by a custom-made LabVIEW software and was delivered through a D/A converter (National Instruments Inc., US). Each stimulus was delivered via a speaker (MSP3, YAMAHA, inc. Japan) which put 1.0 m in front of the participants. To maintain the distance between the participant’s head and the speaker, we instructed the participants to maintain their head location once we measured the distance.

### Somatosensory stimulus

Vibrotactile stimuli were delivered to the fingertip of the index finger via a piezoelectric actuator (Murata Electronics, 7BB-20-6L0). Each stimulus consisted of 200 Hz sinusoidal vibration with a duration of 0.2 s. We used this frequency because the detection threshold of the vibration stimulus was the lowest in this frequency, which we confirmed in a pilot experiment. The actuator was controlled via a custom-made LabVIEW software through the D/A converter. Each participant was instructed to put the pad of the right index fingertip on the surface of the actuator without volitionally pushing down or manipulating the actuator. Before starting the experiment, we confirmed that the auditory signals arising from the vibrotactile stimuli with these properties were under the auditory detection threshold in all participants.

### Intensity discrimination task

This study used a two-alternative forced choice procedure in an intensity discrimination task. Participants were seated in a piano chair with their right hand on a table. Two sensory stimuli with varying stimulus intensities were delivered to participants every trial. The intensity of the first stimulus in a single trial, termed as standard stimulus, was always set to 20 dB for the auditory stimuli and 10 dB for the somatosensory stimuli above the sensory detection threshold that was assessed prior the testing for each participant. The second stimulus, termed as a comparing stimulus, was delivered at the interstimulus interval of 500 ms with varying stimulus intensities. The differences in the intensity between the standard and comparing stimuli ranged from 0.25 to 4 dB with a step of 0.25 dB. The positive and negative value of the intensity of the comparing stimulus means the larger and smaller intensity of the comparing stimulus than the standard stimulus, respectively. Whether the intensity of the comparing stimulus was the positive or negative value was randomly defined every trial. Participants were asked to answer whether they perceive the intensity of the comparing stimulus was larger or smaller than that of the standard stimulus. The intensity of the comparing stimulus was determined adaptively using a weighted up-down staircase method ^31^. Each correct response decreases the difference in the intensity between the two stimuli by 0.25 dB in the next trial, each incorrect response to an increase by 0.75 dB. The intensity discrimination test consisted of 100 trials. In this test, we used different intensity of the standard stimulus between the two modalities in order to minimize the difference in the intensity discrimination threshold in response to each single sensory stimulus. The Bayesian integration or maximum likelihood models posit that multisensory interactions depend on the accuracy of inference of each sensory stimulus^19^. For example, if the accuracy of inference of sensory stimulus from one sensory modality is much worse than the other, multisensory interactions are less likely to be observed. In the discrimination task, the discrimination threshold in response to each single sensory stimulus corresponds to the accuracy of each sensory inference, and the discrimination threshold depends on the intensity of the standard stimulus^30^. Thus we adjusted the intensity of the standard stimulus to match the discrimination threshold between the auditory and somatosensory modalities. In our pilot experiment, we confirmed that the discrimination threshold did not differ between the two modalities when we used the abovementioned intensities of the standard stimulus.

### Data analysis

To obtain a psychometric function of the intensity discrimination performance, we fitted the data obtained from the intensity discrimination task into the cumulative Gaussian distribution function, which defined as follows;

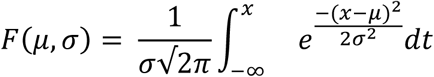

where μ represents how much the center of the function leaves from 0, and σ means the standard deviation of the Gaussian distribution, which is identical to the slope of the function. Because the intensity discrimination task we used was an adaptive procedure, the number of trials differed between intensities of the comparing stimuli. Therefore, we fitted the data by using a weighted non-linear least square algorithm weighted by a square of the number of trials in each intensity. This means that the impact of data with the larger number of trials on the fitting is stronger. The sigma value represents sensitivity of perceiving differences in two stimulus intensities and is defined as threshold. Thus, the present study focused on the sigma value in each condition. We excluded one pianist and one nonmusician from data analyses because they always answered “the intensity of comparing stimulus was larger than that of standard stimulus” if they missed discriminating the intensities of two stimuli. Such a biased answer shifts the center of the psychometric curve abnormally and then impairs accurate estimation of the sigma value.

Sample size was determined so as to fulfill 90% power and a two-tailed 5% level in detecting an interaction between the group and condition factors with an effect size of 0.33, based on our pilot experiment. Statistical analyses were performed using JASP software (JASP Team 2020) and R. A two-way mixed-design analysis of variance (ANOVA) was used to evaluate the sigma and mu values (group × condition). Mauchly’s test was used to assess sphericity before running each ANOVA. The Greenhouse–Geisser correction was performed for nonspherical data. A partial eta squared value (η_p_^2^) was calculated as the effect size for ANOVA. In addition to these analyses, we calculated a Bayes factor (BF_10_) for each analysis to quantify the ratio of the likelihood of an alternative hypothesis to the likelihood of a null hypothesis. For instance, a BF_10_ value of 5 indicates that the alternative hypothesis is 5 times more likely than the null hypothesis, whereas a BF_10_ value of 0.2 indicates that the null model is 5 times more likely than the alternative hypothesis. BF_10_ values above 1 indicate anecdotal, above 3 moderate, and above 10 strong evidence for the alternative hypothesis ^32,33^.

## Results

### Discrimination perception

Figure 1A and 1B shows typical results obtained from the intensity discrimination tasks for one representative pianist and nonmusician, respectively. We fitted the data into the cumulative Gaussian distribution function and obtained the sigma and the mu values that represent the slope and the center of the function, respectively. The dashed line indicates the curve fitted by the psychometric function and the size of the circle indicates the number of trials.

**Figure 1.**
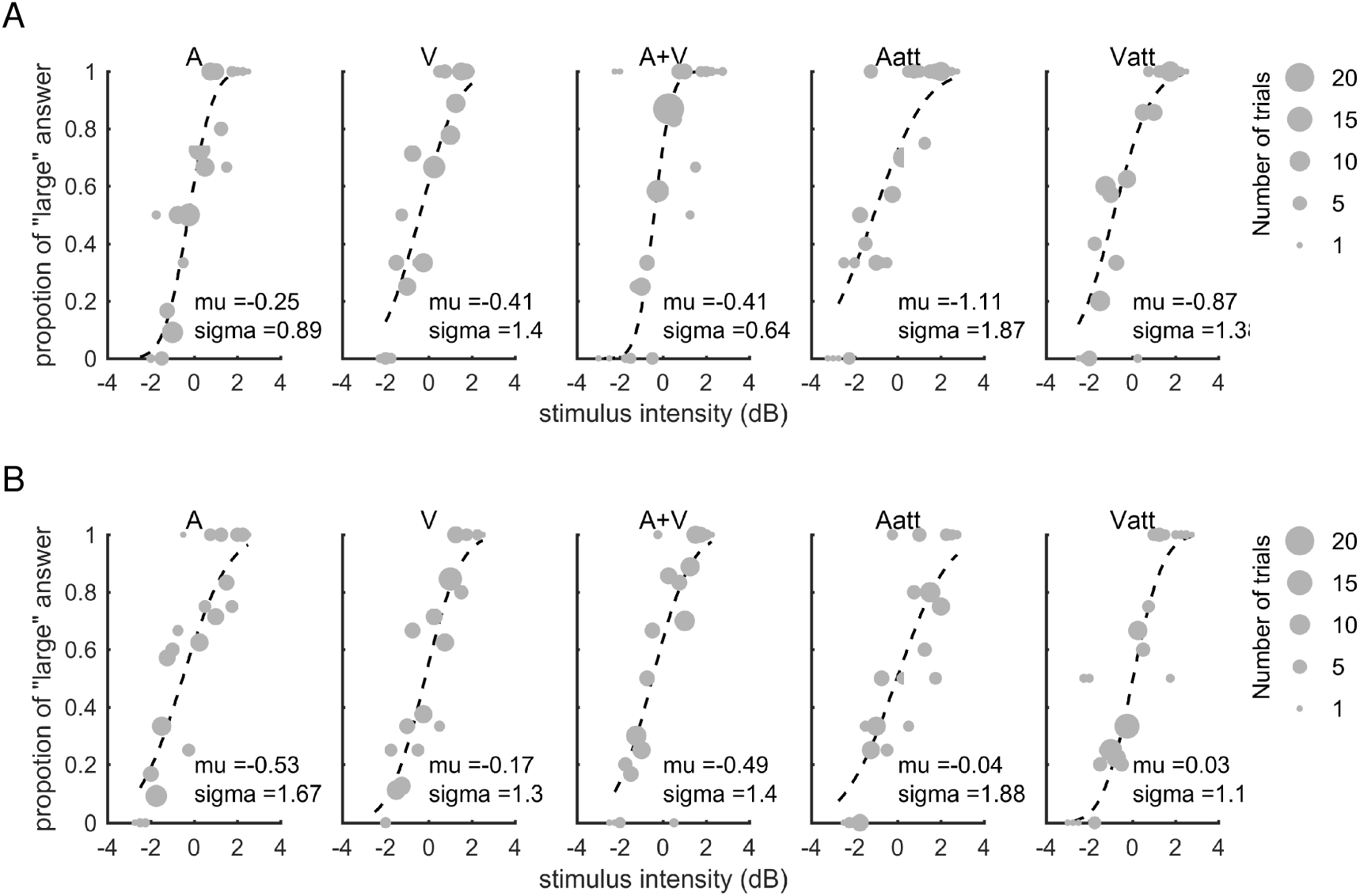
Intensity discrimination performance in unimodal and multisensory conditions. A and B: The panels show the psychometric curves obtained from the intensity discrimination task with the five conditions in a representative pianist (Fig. 1A) and a nonmusician (Fig. 1B), respectively. The dashed line indicates the fitted cumulative Gaussian distribution function. The size of each circle represents the number of trials. The vertical axis means the proportion of answering “the intensity of comparing stimulus was larger than that of standard stimulus” at each stimulus intensity defined as a difference in the intensity between the two stimuli.

### Selective attention

#### Auditory modality

To examine a group difference in an effect of the task-irrelevant vibrotactile stimuli on the auditory discrimination perception, we compared the mu and sigma values obtained from the A and Aatt conditions between the pianists and nonmusicians. Two-way mixed ANOVA revealed no significant effects of the group (F_1,26_=0.69, p=0.42, η_p_^2^=0.03) and condition (F_1,26_=0.52, p=0.48, η_p_^2^=0.02) factors nor the interaction between the two factors (F_1,26_=2.65, p=0.12, η_p_^2^=0.09) on the mu values (supplementary fig. 1A). A Bayesian two-way mixed ANOVA also supports the null hypothesis (supplementary table 1). By contrast, a two-way mixed ANOVA yielded the significant interaction between the group and condition factors on the sigma value (Fig. 2A; F_1,26_=5.25, p=0.03, η_p_^2^=0.17). A Bayesian two-way mixed ANOVA also provided very strong evidence for the group + condition + group×condition model on the sigma values (BF_10_=123.75, table 1). A simple effects test for the interaction further revealed a significant difference in the sigma value in the A condition between the two groups (F_1,26_=4.54, p=0.04, η_p_^2^=0.15, BF_10_=1.80). In addition, the simple effects test revealed a significant difference in the sigma value between the A and Aatt conditions in the pianists (F_1,13_=25.27, p<0.01, η_p_^2^=0.66, BF_10_=138.29), but not the nonmusicians (F_1,13_=2.18, p=0.16, η_p_^2^=0.14, BF_10_=0.66). These results indicate that the unimodal auditory discrimination perception and the effect of the task-irrelevant vibrotactile stimuli on the auditory discrimination perception differed between the two groups.

**Table 1.** Bayesian Repeated Measures ANOVA for the sigma values in auditory selective attention

| **Models** | **P(M)** | **P(M\|data)** | **BF<sub>M</sub>** | **BF<sub>10</sub>** | **error %** |
| --- | --- | --- | --- | --- | --- |
| Null model (incl. subject) | 0.200 | 0.004 | 0.015 | 1.000 |  |
| Condition + Group + Condition * Group | 0.200 | 0.466 | 3.491 | 123.750 | 2.198 |
| Condition | 0.200 | 0.294 | 1.665 | 78.056 | 1.604 |
| Condition + Group | 0.200 | 0.233 | 1.217 | 61.960 | 2.133 |
| Group | 0.200 | 0.003 | 0.012 | 0.783 | 1.350 |

*Note.* All models include subject as a random effect

**Figure 2.**
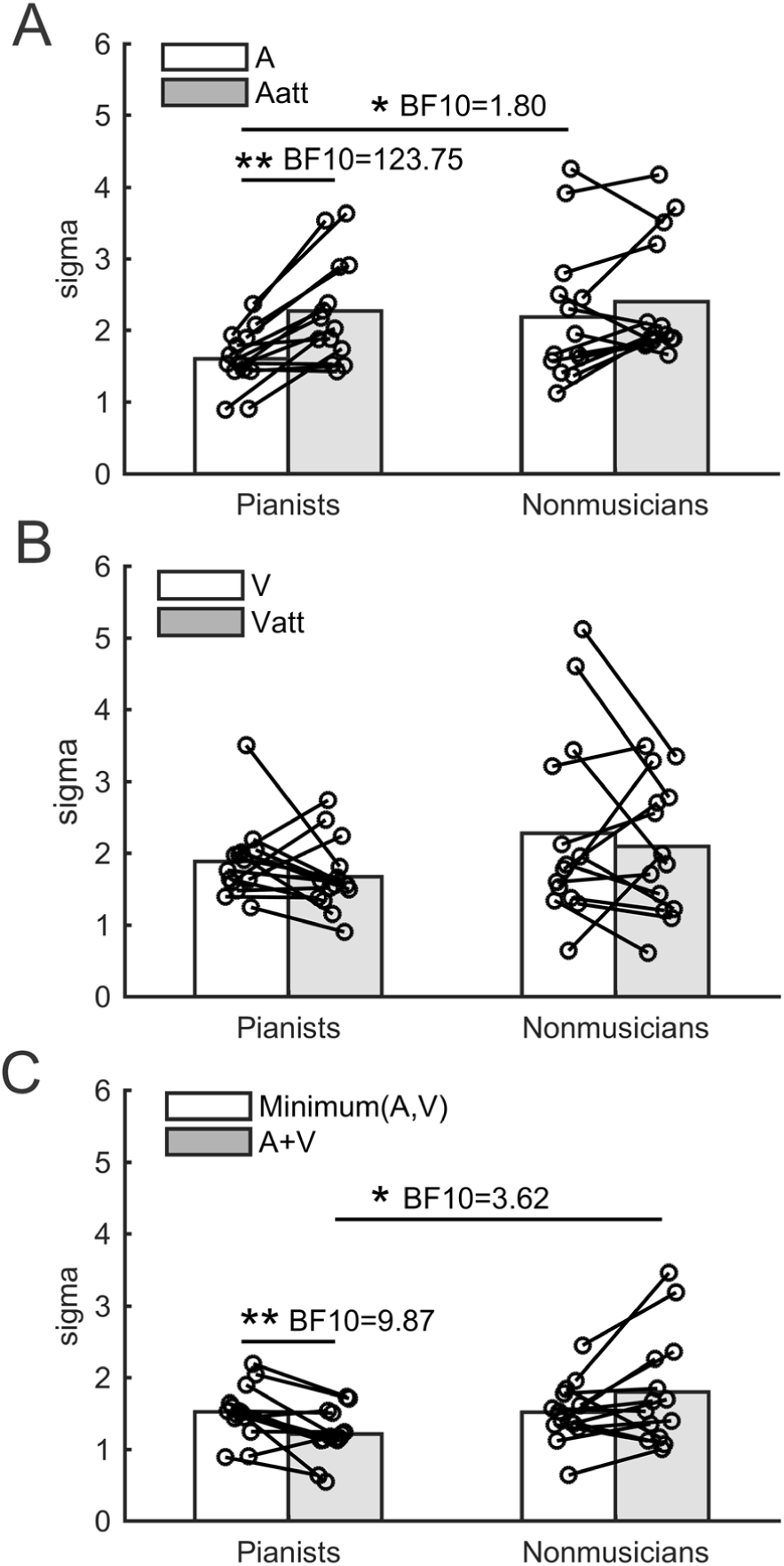
Group mean of the sigma value obtained from each condition in the pianists and nonmusicians. A: The unimodal auditory (A) vs auditory selective attention (Aatt) conditions. B: The unimodal vibrotactile (V) vs vibrotactile selective attention (Vatt) conditions. C: The multisensory integration (A+V) vs the minimum sigma value of those obtained from the A and V conditions (minimum(A,V)). Each circle represents individual data. *,**: p<0.05, 0.01.

#### Vibrotactile modality

To examine a group difference in an effect of the task-irrelevant auditory stimuli on the vibrotactile discrimination perception, we compared the mu and sigma values obtained from the V and Vatt conditions between the pianists and nonmusicians. Two-way mixed ANOVA revealed no significant effects of the group (F_1,26_=0.35, p=0.56, η_p_^2^=0.01) and condition (F_1,26_=1.83, p=0.19, η_p_^2^=0.07) factors nor the interaction between the two factors (F_1,26_=2.21, p=0.15, η_p_^2^=0.08) on the mu values (supplementary fig. 1B). A Bayesian two-way mixed ANOVA also supports the null hypothesis (supplementary table 2). Similarly, a two-way mixed ANOVA yielded no significant main effects of the group (F_1,26_=1.97, p=0.17, η_p_^2^=0.07) and condition factors (F_1,26_=1.30, p=0.27, η_p_^2^=0.05) and their interaction (F_1,26_<0.01, p=0.95, η_p_^2^<0.01) on the sigma value (Fig. 2B). A Bayesian two-way mixed ANOVA also support the null hypothesis (table 2). These results indicate that the vibrotactile discrimination perception did not differ between the two groups and the task-irrelevant auditory stimuli did not affect the vibrotactile discrimination perception in both groups.

**Table 2.** Bayesian Repeated Measures ANOVA for the sigma values in vibrotactile selective attention

| **Models** | **P(M)** | **P(M\|data)** | **BF<sub>M</sub>** | **BF<sub>10</sub>** | **error %** |
| --- | --- | --- | --- | --- | --- |
| Null model (incl. subject) | 0.200 | 0.368 | 2.324 | 1.000 |  |
| Group | 0.200 | 0.285 | 1.594 | 0.775 | 1.297 |
| Condition | 0.200 | 0.169 | 0.812 | 0.459 | 1.423 |
| Condition + Group | 0.200 | 0.133 | 0.614 | 0.362 | 1.456 |
| Condition + Group + Condition * Group | 0.200 | 0.046 | 0.192 | 0.124 | 1.724 |

*Note.* All models include subject as a random effect

### Multisensory integration

To examine a group difference in the multisensory integration, we compared the sigma value obtained from the A+V condition and the minimum sigma value of those obtained from the A and V conditions. Eight pianists and 7 nonmusicians showed better performance (i.e. lower sigma value) in the auditory condition than the vibrotactile condition. For the mu value in the A+V condition and in either the A or V conditions corresponding to the minimum sigma value (supplementary fig. 1C), a two-way mixed ANOVA yielded no significant main effects of the group (F_1,26_=2.34, p=0.14, η_p_^2^=0.08) and condition (F_1,26_=0.33, p=0.57, η_p_^2^=0.01) factors nor the interaction between the two factors (F_1,26_=0.35, p=0.56, η_p_^2^=0.01). A Bayesian two-way mixed ANOVA also supports the null hypothesis (supplementary table 3). For the sigma value, a two-way mixed ANOVA yielded a significant interaction between the group and condition factors on the sigma value (F_1,26_=11.29, p<0.01, η_p_^2^=0.30). A Bayesian two-way mixed ANOVA indicated moderate evidence for the alternative hypothesis (table 3, BF_10_=4.98 for group + condition + group×condition model). A simple effects test for the interaction revealed a significant difference in the sigma value obtained from the A+V condition between the two groups (F_1,26_=6.65, p=0.02, η_p_^2^=0.20, BF_10_=3.62). Furthermore, the simple effects test further revealed significant differences in the sigma value between the A+V condition and the minimum sigma value of those obtained from the A and V conditions in the both groups. In the pianists, the sigma value of the A+V condition was lower than that of the minimum sigma value (F_1,13_=11.29, p<0.01, η_p_^2^=0.46, BF_10_=9.87). By contrast, there was no significant difference in the sigma value between the A+V condition and the minimum one of those obtained from the A and V conditions in the nonmusicians (F_1,13_=3.47, p=0.09, η_p_^2^=0.21, BF_10_=1.05). These results indicate that the simultaneous presentation of auditory and vibrotactile stimuli improved the discrimination perception about 20% in the pianists, but not in the nonmusicians.

**Table 3.**
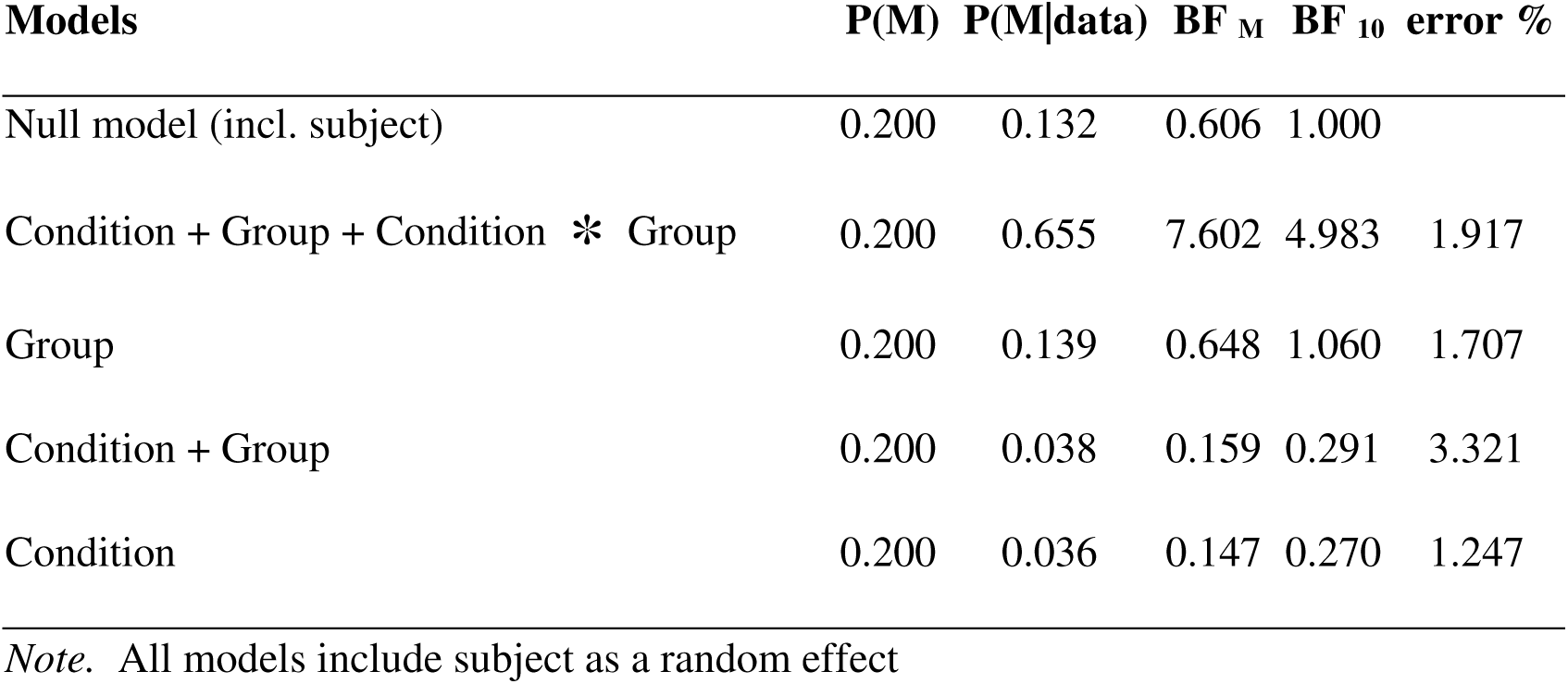
Bayesian Repeated Measures ANOVA for the sigma values in multisensory integration

| **Models** | **P(M)** | **P(M\|data)** | **BF<sub>M</sub>** | **BF<sub>10</sub>** | **error %** |
| --- | --- | --- | --- | --- | --- |
| Null model (incl. subject) | 0.200 | 0.132 | 0.606 | 1.000 |  |
| Condition + Group + Condition * Group | 0.200 | 0.655 | 7.602 | 4.983 | 1.917 |
| Group | 0.200 | 0.139 | 0.648 | 1.060 | 1.707 |
| Condition + Group | 0.200 | 0.038 | 0.159 | 0.291 | 3.321 |
| Condition | 0.200 | 0.036 | 0.147 | 0.270 | 1.247 |

*Note.* All models include subject as a random effect

### Single modality

To examine the differences in the sigma and mu values between the auditory and vibrotactile modalities, we performed a one-way ANOVA on the two values. For the sigma values, we found no significant main effect of the modality (F_1,27_=0.61, p=0.44, η_p_^2^=0.02). A Bayesian one-way ANOVA also support the null hypothesis for the modality effect (BF_10_=0.36) on the sigma values. For the mu values, there were no significant main effect of the modality (F_1,27_=0.22, p=0.65, η_p_^2^<0.01, BF_10_=0.30). These results indicate that the discrimination perception in response to single sensory stimuli did not differ between the two modalities in this experimental setting.

## Discussion

The present study examined multisensory sensory interaction functions at the auditory and somatosensory modalities in pianists and musically untrained individuals. The psychophysical experiments revealed differences in the unimodal and multisensory functions of these modalities between the two groups. The perceptual functions of the unimodal intensity discrimination task were superior in the pianists to the nonmusicians with respect to the auditory modality but not to the somatosensory modality. For the selective attention, the task-irrelevant vibrotactile stimuli interfered the auditory discrimination perception in the pianists, but not the nonmusicians. By contrast, the discrimination perception was improved only in the pianists when the task-relevant auditory and vibrotactile stimuli were simultaneously presented. These findings provide first evidence of unique multisensory interaction functions in highly trained individuals.

### Multisensory integration function

A novel finding of the present study was superior intensity discrimination in the pianists to the nonmusicians when the multisensory stimulus was presented. Several previous studies also demonstrated that musicians could react faster ^34^ and perceive the frequency more accurately ^35^ to multisensory stimulus than nonmusicians, suggesting superior multisensory integration function in musicians to nonmusicians. While these studies compared the reaction time and the frequency discrimination perception between the groups when multisensory stimuli were given, none of them compared those behavioral measures between the multisensory condition and the better one of the individual unimodal conditions. Thus, it remained unclear whether the superior behavioral performance in the multisensory condition in musicians resulted from a superior multisensory integration function or merely reflected the superior perception of each unimodal condition. To resolve this critical issue, the present study investigated both unimodal and multisensory functions and supported the former case. Only in the pianists, the sigma value was lower (i.e. more sensitive) in the A+V condition than the better one of the two unimodal conditions. In piano performance, a weaker keystroke induces smaller intensity of sound and somatosensory feedback, and vice versa. Thus, pianists have abundant experiences of receiving such “matched” multisensory stimuli since childhood, which may develop the specialized multisensory integration function in the intensity domain between the auditory and vibrotactile modalities. In the nonmusicians, by contrast, no significant difference in the discrimination perception between the A+V condition and the better one of the two unimodal conditions suggests that the nonmusicians formed their perception based on information from one of the two modalities, rather than integrating the multisensory information.

### Selective attention in pianists and nonmusicians

We also found that the task-irrelevant auditory stimuli did not affect the vibrotactile discrimination perception in both groups. In contrast, the task-irrelevant vibrotactile stimuli worsened the auditory discrimination perception more in the pianists than nonmusicians, indicating that the function of selectively ignoring the somatosensory information during auditory processing was deteriorated specifically in expert pianists. In the present study, the participants were instructed to ignore the task-irrelevant modality information, which is likely to involve top-down selective attention (suppression of task-irrelevant information and strengthening of task-relevant information) ^36–38^. Our results therefore suggest degradation of a function responsible for suppression of somatosensory information through long-term piano training. Previous studies demonstrated that auditory perturbation applied during performing finger or singing movements less affected control of those movements in musicians than nonmusicians ^18,39,40^, suggesting the improvement of suppressing auditory information during movements in musicians. One of these studies ^39^ further reported that the pianists struck strongly immediately after the auditory perturbation, in order to volitionally facilitate somatosensory information for compensating the disturbed auditory feedback. In fact, musicians do not always play instruments in the same acoustic environment, but play music using instruments with different acoustic properties at various concert halls. Somatosensory information would thus play essential roles in successful performance because of its robustness relative to the auditory information that varies largely depending on the environments. Supporting evidence was that disruption of somatosensory but not auditory information declined singing performance of expert singers ^41,42^, suggesting that expert singers rely more on somatosensory information than on auditory information during singing. It is thus possible that the somatosensory system in musicians has been reorganized through the long-term musical training in order to efficiently process somatosensory information. Such plastic adaptation would shape the robust somatosensory processing but instead compromise inhibition of somatosensory information during auditory processing (i.e., auditory selective attention). This view is comparable to speech, in which somatosensory information also plays an important role in both production and perception of speech ^43,44^.

### Intensity discrimination in unimodal modalities

In line with several previous studies that demonstrated the superior auditory functions in musicians to nonmusicians ^45–49^, we found that, in the intensity discrimination at the auditory modality, the sigma value of the psychometric curve was lower in the pianists than the nonmusicians. Fine perception of the sound intensity during listening and playing musical pieces is an essential skill for crystalizing piano performance. Furthermore, pianists have been exposed to abundant sound from the piano, which suggests that everyday practice induced training-dependent and/or use-dependent plasticity in the auditory functions, and thereby shaped the accurate perception of discriminating sound intensities ^50,51^. By contrast, the intensity discrimination in the somatosensory modality did not differ between the two groups. Contrary to this, previous studies demonstrated superior somatosensory perceptions assessed by a two-point discrimination task and a tactile frequency discrimination task in musicians compared with nonmusicians ^24,35,52,53^. These contrasting results suggest that neural mechanisms underlying the somatosensory discrimination perception differ between the frequency and intensity domains. A discrimination task corresponding to the former domain uses static stimulation to a fingertip to which the slowly adapting mechanoreceptors may respond, whereas one corresponding to the latter domain uses the vibrotactile stimulation same as the present study, which activates fast adapting mechanoreceptors. Although the intensity and frequency discrimination tasks use the vibrotactile stimulation with similar properties, different neural processes in the nervous system mediate these two types of stimuli ^54^. Thus, it is possible that the effects of extensive piano training on somatosensory discrimination perception differ between the two domains. Indeed, a previous study demonstrated no difference in heaviness discrimination perception between pianists and nonmusicians ^52^.

## Limitations

This study used a pure tone for the auditory stimulus, not a complex piano sound. Previous studies demonstrated that auditory-tactile interaction is more pronounced when the auditory stimulus consists of a complex tone than of a pure tone^28,55^. This suggests that the differences in the auditory-somatosensory interactions between the pianists and non-musicians found in this study are more emphasized if piano tone is used as the auditory stimulus. However, the observed differences in the auditory-somatosensory interactions between the two groups, even with pure tone stimuli, indicate the remarkable differences in the interactions between the two groups.

## Conclusion

The present study examined the multisensory interaction functions in trained individuals. The results from the psychophysical experiments firstly demonstrated that highly trained pianists were superior in the multisensory integration function and inferior in the robustness of the auditory processing against task-irrelevant somatosensory stimuli compared with those of nonmusicians. The extensive auditory-somatosensory experiences through daily piano practicing would shape the unique multisensory interaction functions, which enables pianists to meaningfully integrate the auditory and somatosensory information but instead exacerbates the top-down selective inhibition of somatosensory information during auditory processing.

## Funding

This work was supported by Grant-in-Aid for JSPS Fellows (to M.H.) and CREST (Core Research for Evolutional Science and Technology) of the Japan Science and Technology Agency (to S.F.).

## DISCLOSURES

No conflicts of interest, financial or otherwise, are declared by the authors.

## Notes

### Competing Interest Statement

The authors have declared no competing interest.

