## Supplemental Tables for "Multisensory interactions on auditory and somatosensory information in expert pianists"

| Supplementary table 1. Bayesian Repeated Measures ANOVA for the mu value in auditory selective attention | | | | | | | | | | |
| --- | --- | --- | --- | --- | --- | --- | --- | --- | --- | --- |
| **Models** | | **P(M)** | | **P(M\|data)** | | **BF _M_** | | **BF _10_** | | **error %** |
| Null model (incl. subject) |  | 0.200 |  | 0.474 |  | 3.609 |  | 1.000 |  |  |
| Group |  | 0.200 |  | 0.224 |  | 1.156 |  | 0.473 |  | 1.412 |
| Condition |  | 0.200 |  | 0.155 |  | 0.732 |  | 0.326 |  | 1.497 |
| Condition + Group |  | 0.200 |  | 0.074 |  | 0.322 |  | 0.157 |  | 1.258 |
| Condition + Group + Condition  ✻  Group |  | 0.200 |  | 0.072 |  | 0.312 |  | 0.153 |  | 1.639 |
| *Note.*  All models include subject | | | | | | | | | | |

| Supplementary table 2. Bayesian Repeated Measures ANOVA for the mu value in vibrotactile selective attention | | | | | | | | | | |
| --- | --- | --- | --- | --- | --- | --- | --- | --- | --- | --- |
| **Models** | | **P(M)** | | **P(M\|data)** | | **BF _M_** | | **BF _10_** | | **error %** |
| Null model (incl. subject) |  | 0.200 |  | 0.396 |  | 2.619 |  | 1.000 |  |  |
| Condition |  | 0.200 |  | 0.213 |  | 1.085 |  | 0.539 |  | 1.527 |
| Group |  | 0.200 |  | 0.198 |  | 0.989 |  | 0.501 |  | 0.929 |
| Condition + Group |  | 0.200 |  | 0.107 |  | 0.482 |  | 0.272 |  | 1.209 |
| Condition + Group + Condition  ✻  Group |  | 0.200 |  | 0.085 |  | 0.373 |  | 0.215 |  | 2.099 |
| *Note.*  All models include subject | | | | | | | | | | |

| Supplementary table 3. Bayesian Repeated Measures ANOVA for the mu value in A+V condition | | | | | | | | | | |
| --- | --- | --- | --- | --- | --- | --- | --- | --- | --- | --- |
| **Models** | | **P(M)** | | **P(M\|data)** | | **BF _M_** | | **BF _10_** | | **error %** |
| Null model (incl. subject) |  | 0.200 |  | 0.373 |  | 2.383 |  | 1.000 |  |  |
| Group |  | 0.200 |  | 0.297 |  | 1.694 |  | 0.797 |  | 4.661 |
| Condition + Group + Condition  ✻  Group |  | 0.200 |  | 0.126 |  | 0.577 |  | 0.338 |  | 72.039 |
| Condition |  | 0.200 |  | 0.116 |  | 0.525 |  | 0.311 |  | 1.692 |
| Condition + Group |  | 0.200 |  | 0.087 |  | 0.381 |  | 0.233 |  | 1.105 |
| *Note.*  All models include subject | | | | | | | | | | |
